# Spatiotemporal analysis of F-actin polymerization with micropillar arrays reveals synchronization between adhesion sites

**DOI:** 10.1101/2024.06.22.600020

**Authors:** Sarit Hollander, Yuanning Guo, Haguy Wolfenson, Assaf Zaritsky

## Abstract

We repurposed micropillar-arrays to quantify spatiotemporal inter-adhesion communication. Following the observation that integrin adhesions formed around pillar tops we relied on the precise repetitive spatial control of the pillars to reliably monitor F-actin dynamics in mouse embryonic fibroblasts as a model for spatiotemporal adhesion-related intracellular signaling. Using correlation-based analyses we revealed localized information-flows propagating between adjacent pillars that were integrated over space and time to synchronize the adhesion dynamics within the entire cell. Probing the mechanical regulation, we discovered that stiffer pillars or partial actomyosin contractility inhibition enhances inter-adhesion F-actin synchronization. Our results suggest that adhesions can communicate and highlight the potential of using micropillar arrays as a tool to measure spatiotemporal intracellular signaling.

## Introduction

Cells coordinate their structure and signaling in space and time to mediate diverse functions such as spreading, migration and division. Mechanistic investigations of how signaling leads to function are enabled by using fluorescent markers for quantification of spatiotemporal signaling patterns, analyzing their relationships with the cell’s morphodynamics, and how they alter under different experimental conditions. Several examples include revealing the concept of actin retrograde flow from the cell’s leading edge toward the cell’s interior and its involvement in migration [1], Rac1-mediated protrusion waves [2], Rho GTPase coordination during cell protrusion [3], Asef (GEF) regulation of Cdc42 and Rac1 (GTPases) control of cell edge dynamics [4], PI3K-mediated [5], CDC42-mediated [6] or Ca2+-mediated [7] alteration of protrusions to reorient cell polarity during migration, and deciphering the regulation of actin nucleators in lamellipodia formation [8, 9].

Integrin-mediated cellular interactions with the extracellular matrix (ECM) are fundamental and essential for numerous cellular functions, including cell migration, survival, and proliferation [10], that are dependent on cellular anchorage to the environment. Beyond serving as anchoring sites, integrin adhesions are signaling centers that recruit key signaling proteins to the cytosolic side of the adhesions, including kinases such as src or focal adhesion kinase (FAK), and rho GTPases [11, 12]. Moreover, ion channels that interact or are present in the vicinity of integrin adhesions (e.g., piezo channels) can give rise to local flow of calcium ions that can regulate the adhesions themselves via calcium-dependent enzymatic activity (e.g., calpain-dependent cleavage of FAK and talin) or the actin cytoskeleton that (indirectly) associates with the integrins (e.g., through sequestration of the profilin-G-actin complex [13]). Given that many of the signaling molecules can dissociate from the adhesions and undergo rapid diffusion in the cytoplasm, and given the mechanosensitivity of piezo channels which can enhance calcium flow upon adhesion growth [14], an intriguing possibility is that neighboring adhesions can communicate through the activity of those molecules/ions, thereby leading to local synchronization in activation of signaling pathways between adhesions.

Most current methods for quantification of spatiotemporal signaling rely on local measurements of fluorescent intensities followed by temporal alignment and averaging to generate shape-invariant spatiotemporal cell representations. This is commonly achieved by extracting the protrusion and signaling dynamics along the cell edge’s by partitioning the cell boundary to many “quantification windows”, tracking these windows, and recording the corresponding subcellular time-series [2]. However, the ability to reliably track interior windows located further away from the cell’s edge is severely limited due to the accumulation of tracking errors caused by cell movement and shape deformations. To tackle this challenge of reliably monitoring signaling across the entire cell, Jiang et al. proposed using a separate fluorescence channel (e.g., VASP - punctate, profilin - diffusive) as a location fiducial, matching fiducial images to a reference frame using nonlinear registration, and using this registration and optical flow to extract signaling (actin) time series at every subcellular location [15]. While showing effective spatiotemporal quantification, this approach is still sensitive to the ability to reliably register the fiducial channel as the cell undergoes complex deformation patterns.

Here we propose to repurpose a micropillar-array system, originally designed to measure traction forces [16, 17, 18], as location fiducials for adhesion-related intracellular signaling. The micropillars are coated with an ECM protein and are tightly spaced such that cells remain on top of the pillars. Each pillar constitutes an independent site for integrin adhesions to form [18, 19] enabling reliable extraction of time series in relation to each pillar that is not sensitive to cell deformations or to alterations of fluorescent fiducial channels. The micropillar-array can be thought of as fiducials that provide a systematic and stable repetitive template that maps the entire cell in a periodic pattern with predefined resolution. This precise spatial control can enable systematic investigation of inter-adhesion communication. We applied this micropillar-array system to investigate inter-adhesion communication in mouse embryonic fibroblasts. We tracked filamentous actin (F-actin) dynamics since F-actin interacts with integrin adhesions (e.g. via talin) and plays a major role in regulating adhesion dynamics, and since F-actin can be triggered to locally “turn on” (namely, polymerization initiation) through cytosolic signals, e.g. via formins [17]. Correlation analysis confirmed that long-range intracellular synchronization is facilitated through integration of localized information flows propagating between adhesions on adjacent pillars. Probing the mechanical regulation, we revealed that stiffer substrate and partial actomyosin contractility inhibition both induce enhanced inter-adhesion F-actin synchronization. These results indicate the existence of inter-adhesion communication and suggest the potential of using micropillar arrays as a tool to measure spatiotemporal intracellular signaling.

## Results

### Repurposing micropillars arrays to measure intracellular synchronization and information flow

We repurposed the micropillar-array system, initially designed for measuring cellular forces, to serve as landmarks for tracking spatiotemporal intracellular signaling. We acquired 81 2D time-lapse confocal microscopy imaging of 13 mouse embryonic fibroblasts (MEFs) expressing tdTomato-tractin (a marker for filamentous actin, F-actin) [20] positioned on top of an array of micropillars, each 5.3-μm high and 2-μm wide, with a 4-μm distance between any two adjacent pillars. The cells formed adhesions that assembled around the micropillars (Fig. S1), leading to local accumulation of F-actin around the pillars, which were monitored for approximately 60 minutes (Fig. S2) with temporal resolution of ∼30 seconds between frames (Fig. 1A, <u>Video S1</u>).

**Figure 1.**
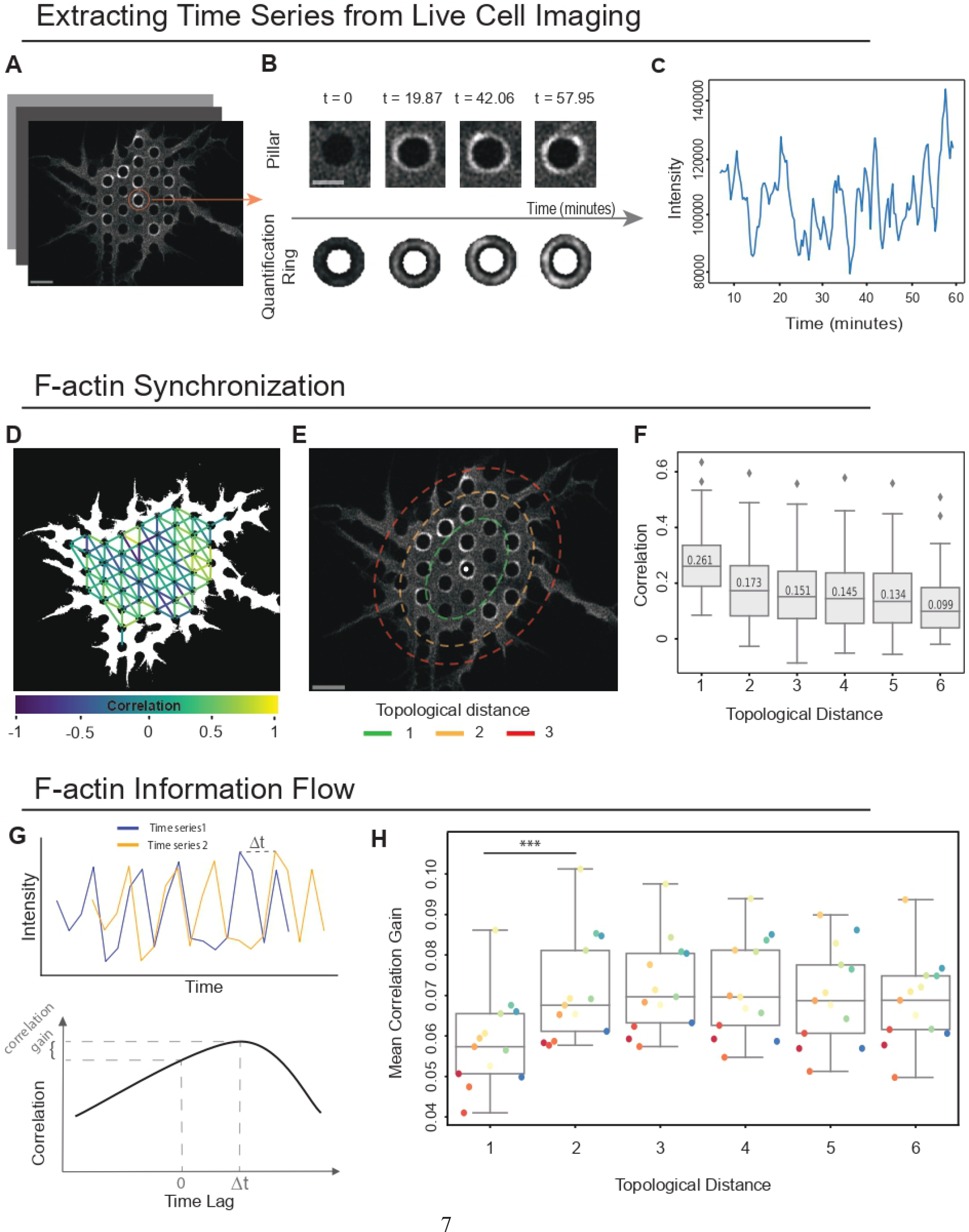
Repurposing micropillar arrays to measure local synchronization. **(A-C)** Schematic sketch of measuring intracellular protein dynamics. (**A**) Live imaging of single cells placed on top of the surface of a micropillars array (black circular voids). Scale bar = 5 m. **(B)** A pillar (top) and its corresponding quantification ring (bottom) over time. Scale bar = 2 m. **(C)** The time series extracted from the mean intensity in a pillar’s quantification ring over time. **(D)** Visualization of the correlation between the adjacent pillars’ time-series. Edges connect adjacent pillars and are colored according to their correlation. **(E)** Topological distance is calculated according to the 8-pillar neighborhood around each pillar. Shown are topological distances of 1 (green), 2 (yellow), and 3 (red) in respect to the middle pillar (white dot). Note that the 8-pillar neighborhood defines an oval-like topological distance. Scale bar = 5 m. **(F)** The mean correlation of all pillar pairs pooled across each cell as a function of the pairwise topological distance. Number of cells at each topological distance (N) and the mean temporal correlation between the pillar pairs time series (R): N1 = 81, R1 = 0.27, N2 = 81, R2 = 0.19, N3 = 814, R3 = 0.17, N4 = 81, R44 = 0.16, N5 = 74, R5 = 0.15, N6 = 41 R6 = 0.12. **(G)** Depiction of calculating the cross correlation between two time series. Top: time series #2 is temporally shifted by Δt relative to time series #1. Bottom: correlation gained by the shifted time series. **(H)** The per cell average correlation gain for all pillar pairs according to their topological distance with a maximum time lag of 3 frames (± 90 seconds). Each data point represents a cell. Each cell is color coded. The mean correlation gain between topological distance 1 to topological distance 2 was deemed statistically significant using the Wilcoxon signed-rank statistical test rejecting the null hypothesis that the difference in correlation gain was distributed around zero (N = 13 cells, *** - p-value < 0.001).

We segmented and tracked the pillars over time, and quantified the tdTomato-tractin fluorescence intensity in a ring of width 1.1 μm surrounding each pillar, where an adhesion-induced enrichment in F-actin was observed (Fig. 1B). The fluorescence intensity over time in each quantification ring defined the F-actin time series associated with each pillar (Fig. 1C). To enable quantitative comparison across experiments, we first subtracted the background, defined for each time frame according to the mean tdTomato-tractin intensity within all pillars at that time (Fig. S3). Next, we normalized each quantification ring’s time series against its own mean and standard deviation using z-score normalization. This normalization scales the fluorescent signal in each time frame as the number of standard deviations away from that pillar’s mean across time.

To determine the local intracellular (i.e., pillar-to-pillar) F-actin synchronization we calculated the correlation between adjacent pairs of pillars’ time-series. Most pairs showed positive correlations (Fig. 1D) much higher than the control pairwise correlations derived from the time-series extracted from the interior of the pillars indicating the existence of inter-adhesion communication (Fig. S4). We assessed the sensitivity of our measurement by changing the location and the width of the quantification ring and showing consistent pairwise temporal correlations between adjacent pillars confirming the robustness of our measurement (Fig. S5). We next measured the temporal correlations between all pairs of pillars grouped by their topological distances and identified a negative correlation between the topological distance between the pillars and the correlation in their time-series (Fig. 1E-F). The decrease in correlation between pairs of pillars as a function of topological distance suggested that long-range intracellular synchronization was mediated by localized pillar-to-pillar information flow induced by communication between adjacent pillars’ adhesions.

Next, we hypothesized that localized pillar-to-pillar information flow could be manifested by a temporal lagged correlation to pillars that are more distant away reflecting the temporal shift that is required for the signal to propagate between distant pillars. To test this hypothesis we measured all pillar pairs correlation gain by cross-correlation with time lags and grouped them according to their topological distances. Specifically, the correlation between two time-series was calculated for time lags ranging from -3 to +3 frames (i.e., ± 90 seconds), and the correlation gain was the deviation of the maximal time-lagged correlation from the correlation without time shift (Fig. 1G). This analysis confirmed increased correlation gain for pillar pairs beyond immediate adjacency, indicating signal propagation in space over time (Fig. 1H). These results suggest that local pillar-to-pillar information flow is integrated across time and space leading to longer ranged information flow and synchronization. From a methodological perspective, these results support the potential of repurposing micropillar arrays to measure spatiotemporal intracellular signaling.

### Mechanical regulation of Actin intracellular synchronization

Substrate stiffness is known to influence cells’ structure, organization and function. With micropillar arrays, the effective stiffness can be modulated by changing the micropillars’ height, which also influences adhesion formation and cell spreading [21]. We hypothesized that changing the pillars’ effective stiffness would also alter the local inter-adhesion synchronization. To test our hypothesis, we collected a second dataset of 72 2D time-lapse confocal microscopy imaging of 13 tdTomato-tractin MEFs positioned on 13.2 μm height micropillars (whose spring constant is ∼2 pN/nm), and compared the localized pillar-to-pillar correlations to the first dataset of cells on 5.3 μm height micropillars (whose spring constant is ∼31 pN/nm). Building on our results showing higher correlations for adjacent pillars, we measured and plotted for each cell the mean correlation of all pairs of adjacent pillars (y-axis), and the corresponding mean correlation of all pairs of non-adjacent pillars (x-axis) (Fig. 2A). A cell above the diagonal y = x (dashed line) indicates that the correlations between adjacent pairs exceeded that of the non-adjacent pairs. The vast majority of time-lapses, 81/81 on 5.3 μm pillars and 60/72 on 13.2 μm pillars, were above the diagonal, indicating that local synchronization can be measured for both substrate stiffnesses. Direct measurement of the difference between the matched correlations of adjacent and nonadjacent pillar pairs showed enhanced local synchronization for the stiffer pillars of height 5.3 μm (Fig. 2A, inset). Altogether, these results indicate enhanced localized intracellular F-actin synchronization probably via inter-adhesion communication on stiffer substrates.

**Figure 2.**
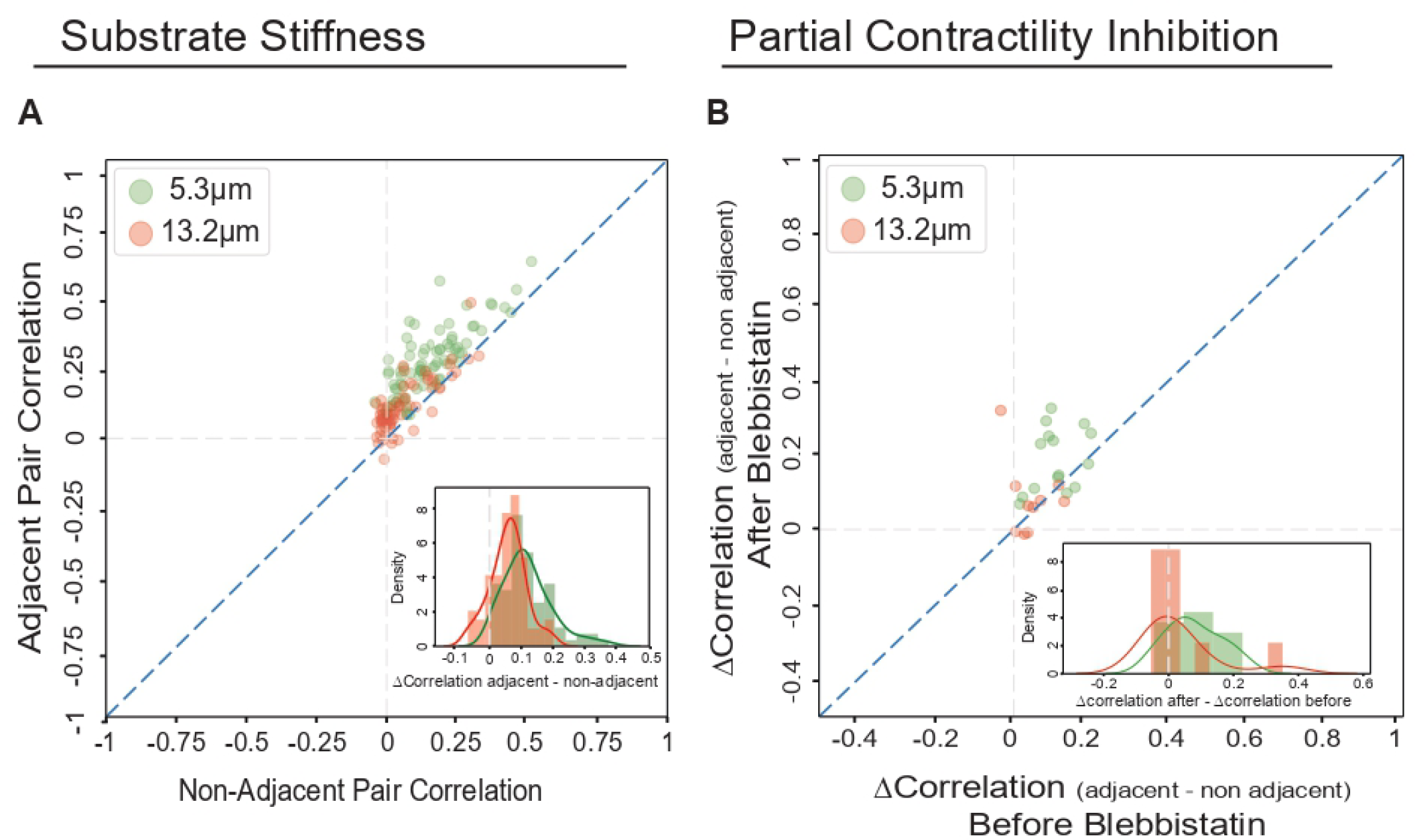
Mechanical regulation of Actin intracellular synchronization. (A) Substrate stiffness. Cells on top of pillars of height 5.3 Dm (N = 81, green) and of height 13.2 Dm (N = 72, red). Each data point records the mean correlation in a movie of a cell’s adjacent (y-axis) and non- adjacent (x-axis) pillar pairs. Adjacent pairs had higher correlations than non-adjacent pairs in all 81 movies on top of pillars of height 5.3 Dm (T-test p-value < 0.0001), and in 83.33% (60/72) of movies on top of pillars of height 13.2 Dm (T-test p-value < 0.001). Inset: Difference between the matched correlations of adjacent and nonadjacent pillar pairs. The stiffer 5.3μm pillars showed enhanced local synchronization in respect to the 13.2μm pillars (T-test p-value < 0.0001). **(B)** Partial contractility inhibition. Matched analysis before (x-axis) and after (y-axis) adding blebbistatin. Each data point records a cell’s change in the correlation of adjacent versus non-adjacent pillar pairs (Δ*correlations*) before and after the treatment. For cells on top of 5.3 Dm height (N = 15, green), the post-blebbistatin Δ*correlations* were higher for 12/15 cells (T-test p-value < 0.01). For cells on top of 13.2 Dm height (N = 10, red), the post-blebbistatin Δ*correlations* were higher for 6/10 cells (p-value not significant). Inset: difference between the matched Δ*correlations* before and after adding blebbistatin. The stiffer 5.3μm pillars (green) showed enhanced local intracellular synchronization in respect to the 13.2μm pillars (red).

Previous studies have shown that slightly decreased myosin II activity can improve intracellular passive forces transmission, and that elevated myosin activity can disrupt these mechanical signals [22, 23, 24]. Thus, we hypothesized that relatively low doses of the myosin II inhibitor Para-nitroblebbistatin (henceforth referred to as ‘blebbistatin’), will enhance the local intracellular synchronization. To assess this hypothesis, we imaged cells on top of 5.3 μm and 13.2 μm micropillars and measured the correlations before versus after 2-3 hours treatment with 5 μM of blebbistatin. Consistent with the aforementioned results, the correlations between adjacent pillar pairs exceeded that of the non-adjacent pairs for both substrate stiffnesses, and both before and after adding blebbistatin, indicative of local synchronization in all conditions (Fig. S6). To compare the change in intracellular synchronization following blebbistatin treatment, we performed a matched analysis where for each cell we measured the change in the correlation of adjacent versus non-adjacent pillar pairs (denoted Δ*correlation*) before and after adding blebbistatin. Plotting the Δcorrelations of cells on top of 5.3 μm pillars, before (x-axis) versus after (y-axis) blebbistatin treatment showed that the post-blebbistatin Δcorrelation was higher (i.e., above the diagonal) for 12/15 cells (p-value < 0.01, Fig. 2B, green). However, blebbistatin did not increase the Δcorrelation of cells on the softer pillars, with only 6/10 cells above the diagonal (Fig. 2B, red). These results suggest that partial contractility inhibition affects F-actin local intracellular synchronization in a rigidity-dependent manner. Specifically, inter- adhesion communication is more susceptible for stiffer substrates.

## Discussion

We propose the repurposing of micropillar-arrays as location fiducials toward quantification of spatiotemporal adhesion-related signaling patterns. Given the formation of focal adhesions around the pillar tops, the micropillar-array defines a patterned arrangement of fiducials that are stably covering the entire cell. The pillars are easy to segment and track, and do not require an additional fluorescent channel, thus facilitating the reliable monitoring of signaling across space and time throughout the entire cell. Micropillar-arrays fiducials overcome limitations of methods that accumulate tracking errors in regions away from the cell edge due to cell motion and deformations [2, 1, 3, 4, 5, 6, 7, 8, 9, 25], or location fiducials methods that require an additional fluorescent channel and may not be homogeneously distributed and include regions that are difficult to track [15]. The size and the distance between pillars can be flexibly optimized and controlled according to the experimental system. Using correlation-based analyses we demonstrated that our method can measure intracellular synchronization and information flow in the context of focal adhesion-mediated communication via monitoring of F-actin dynamics.

Specifically, long-range intracellular synchronization was achieved through integration of localized propagating information flows between adjacent pillars that were enhanced by stiffer substrates and by partial actomyosin contractility inhibition. The evenly spatially distributed pillars can be used in the future to systematically investigate intracellular spatial heterogeneity in adhesion signaling [25], and correlating spatiotemporal adhesion signaling patterns to different cellular compartments with multiplexed fluorescent imaging.

Our analysis relied on inter-adhesion communication measured from the polymerization of F-actin (measured through the local tomato-Tractin density) at focal adhesion sites assembled around the tops of the micropillars. A similar approach can be used to measure spatiotemporal intracellular signaling of focal adhesion complex proteins (e.g., vinculin, paxillin). Such communication can potentially be the source of cellular-level regulation over events that occur in multiple adhesions, perhaps as a feed-forward mechanism to facilitate accumulation of signals at different locations across the cell. In principle, the micropillar-arrays should be able to measure intracellular synchronization and information flow in signaling pathways that do not necessarily interact with the pillars, for example punctate or diffusive molecules. This would be possible by using the pillars similar to other location fiducials methods [15], enjoying the benefits of trackable patterned fiducials that cover the entire cell and change along with the local cell deformations.

A notable limitation of our method is the fixed location of the pillars. Cells exerting forces on the substrate can alter the pillars’ locations, and these changes are tracked in our method. However, migrating or highly deforming cells will change their location with respect to the pillars leading to a pillar time series that reflects a combination of different subcellular time series. This can be partially resolved (in migration) by tracking the cell and adjusting the quantification window.

Realistically, this limitation implies that the optimal use of micropillar-arrays for spatiotemporal signaling quantification is most suitable for short time scales, cells that do not undergo large deformations and non-migrating cells. In addition, the physical size of the pillar is restricting the spatial resolution that can be achieved, with respect to windowing techniques. .

Our findings indicate that cells positioned on stiffer micropillars exhibit enhanced local synchronization, which can play a crucial role in intracellular molecular interaction, communication, and vital signal transduction pathways. This is in line with prior research that connects cell behaviors to variations in pillar height and resulting substrate stiffness [21, 23]. The microenvironment’s rigidity is known to impact focal adhesion initiation and maturation, as well as its downstream mechanosensing and mechanotransduction processes, which leads to different cell behaviors and fates (e.g., proliferate, death, migration, invasion) [26]. Moreover, substrate rigidity dramatically affects the assembly and organization of the actomyosin cytoskeleton that is in contact with the adhesions [23, 27], including F-actin assembly, disassembly, and arrangement [28]. These biomolecular events can be reflected, observed, measured, and evaluated by our micropillar-based platform and method. Of potential importance for the inter-adhesion communication the we observe via F-actin is the recent finding that the activity of the actin elongation factor mDia1 is triggered by actomyosin contractility [29]. Thus, we postulate that enhanced adhesion maturation on stiff matrices, which in turn leads to enhanced transmission of actomyosin forces, leads to high mDia1 activation. This then triggers F-actin polymerization in nearby adhesion sites through the diffusion of activated mDia1 locally. Similar mechanisms are possible through local diffusion of other factors such Ca2+ ions and/or Rho GTPases. These scenarios should be tested in future studies. Moreover, they should be tested also in the context of different levels of contractility inhibition (using different blebbistatin doses) given the higher Δcorrelation before and after blebbistatin treatment on stiff, but not on soft pillars (Fig. 2).

## Methods

### Experimental methods

#### Pillar fabrication and fibronectin coating

To fabricate the pillars, Polydimethylsiloxane (PDMS) was added in a 10:1 ratio to the curing agent (Sylgard 184, Dow Corning) into silicone molds with predefined hole depths and spacings. The molds were then inverted onto glass-bottom 35-mm dishes (D35-20-0-N, Cellvis, Mountain View, CA) and incubated at 60°C for 12 hours to solidify the PDMS. Subsequently, the molds were carefully removed while submerged in 100% ethanol to avoid pillar collapse. Next, the ethanol was removed by replacing it with phosphate-buffered saline (PBS). Human plasma full-length fibronectin (FC010, Merck Darmstadt, Germany) was then applied to the dish at a concentration of 10 μg/μl for an incubation period of 1 hour at 37°C. This step was followed by washing with a PBS buffer to remove fibronectin residuals.

The rigidity of the PDMS pillar arrays, or the external rigidity, was modulated only by altering the height of the pillars because the pillar mold yielded a constant cross-sectional area (2 μm-diameter circle) for each pillar, and because PDMS maintains stable chemical properties.

The spacing from the center of one pillar to the center of the next pillar was set at 4 μm, with pillar heights of 5.3 or 13.2 μm, corresponding to spring constants of 31 and 2 pN/nm, respectively. These values were calculated using Euler-Bernoulli beam theory:

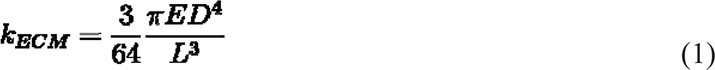

Here, *D* and *L* represent the diameter and height of the pillar, respectively, and *E* is the Young’s modulus of the material. For our experiments, the Young’s modulus of PDMS, (*k_ECM_*), was 2 MPa. By applying a commonly used relation to deduce an effective elastic modulus corresponding to a specific rigidity [30]:

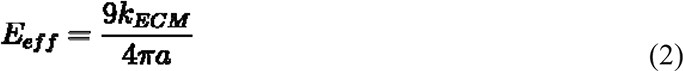

where *a* is the radius of the pillars, we derived effective elastic moduli approximately 1.5 and 22.7 kPa (corresponding to the heights of 13.2 and 5.3 μm, respectively), which falls within the physiologically relevant spectrum, spanning from endothelial tissues to cartilage [31].

#### Cells culture and reagents

To track spatiotemporal intracellular F-actin signaling in living cells, tdTomato-tractin mouse embryonic fibroblasts (MEFs) were used and transfected with pCMV-tdTomato-tractin plasmid, a gift from M. Sheetz (MBI Singapore and the University of Texas Medical Branch) [31].

Briefly, F-tractin is a 43-amino acid peptide fragment in the N-terminus part of ITPKA, which has an F-actin binding ability [20, 32, 33]. tdTomato-tractin not only preserve the actin polymerization and depolymerization rates, but it also binds only to F-actin (without binding to G-actin), thus labeling live cells and allowing tracking of fiber dynamics [20, 32, 33].

tdTomato-tractin MEFs were cultured at 37°C in a 5% CO2 incubator in Dulbecco’s modified Eagle medium (Biological Industries, BI) supplemented with 10% fetal bovine serum (FBS, Gibco) and penicillin-streptomycin (P/S, 100 IU/ml; Biological Industries). During confocal microscopic imaging, MEFs were seeded on micropillar arrays casted on a 35 mm glass-bottom dish, which is kept in a 37°C heating chamber. The medium was replaced by HBSS (BI) with 20 mM HEPES (BI), 10% FBS, and 100 IU/ml P/S.

#### Live imaging

We used a Zeiss LSM800 confocal microscope (20× objective, spatial resolution of 6.386 px/ m) with a heating chamber to preserve temperature of 37°C during live tdTomato-tractin imaging at temporal resolution for 30 seconds per frame. Although Definite Focus was set, there still existed slight focus shifts during live imaging, especially in the beginning of each experiment. To ensure focused imaging, we set the focal plane, halted imaging upon sample displacement out of focus, followed by re-setting the focal plane and imaging. Thus a single cell was discontinuously imaged to generate several time-lapse sequences, the duration distribution of which is shown in Fig. S2.

#### Contractility inhibition experiments

To compare F-actin flow differences and cellular behaviors for the same cells, the nonmuscle myosin II inhibitor Para-nitro blebbistatin (DR-N-111, motorphama) was used to reduce the actomyosin contractility. Cells were treated with 5 μM blebbistatin following 2-3 hours of live imaging. In some cases, cells lost the bright F-actin rings following blebbistatin treatment and were therefore eliminated from the analysis to keep the comparison consistent.

#### Data description

The data used in this study is summarized in <u>Table S1</u>. It included 10-15 cells and 46-82 time- lapse images of these cells per experimental condition.

### Analysis

#### Stage drift correction

To mitigate potential artifacts arising from errors in microscopy re-positioning, we performed a global positioning correction by adjusting the position of each frame relative to the previous one, using the repositioning error calculated with skimage registration function (phase_cross_correlation). The uniformity and predefined grid arrangement of the pillars greatly aided this process, as they served as reliable reference points. These pillars, being identical to each other and systematically organized, provided us with accurate prior information regarding their expected locations.

#### Cell segmentation and pillar tracking

We use a semi-automatic pipeline to segment and track the pillars. We leveraged the pre-defined spatial arrangement to annotate the pillars’ centers of all pillars in the first frame of a video. The pillars maintain their spatial arrangement, but slightly move due to cell motion, cell deformation, and exertion of traction forces on the substrate. To track the pillar exact location in each frame we performed template matching to the pillar interior void (Fig. 1B). Cell deformation and motion could include or exclude new pillars during the experiment. We determined the time intervals for each pillar according to a ratio between the average intensity of the pillar interior void and the ring surrounding it exceeding 0.5. Pillars’ time series durations were determined according to the time the pillar appeared within the cell segmentation mask.

#### Measuring pillars’ F-actin signaling

We monitored F-actin fluorescence fluctuations over time for each pillar using a quantification ring with area of ∼8.13µm 2 formed by two circular masks of different radii (small radius ∼0.63μm, large radius ∼1.73μm, when the pillar’s radius was 1μm). The intensity was integrated across all pixels within the quantification ring (Fig. 1B). The quantification ring’s time series was extracted (Fig. 1C) and normalized against its own mean and standard deviation using z-score normalization. To minimize the effect of background noise on the correlations, the average signal from the pillar interior voids at each time point was subtracted from the time series of both the pillar voids and the original signal in the quantification ring. Pearson correlation was used to measure the correlation between adjacent and non-adjacent pillar pairs. For each cell, we calculated the mean correlation among adjacent pillars and the mean correlation among non-adjacent pillars, and used it in our analysis (Fig. 2A, Fig. 2B). Correlation analysis might be sensitive to ring size: wider rings accumulate more F-actin fluorescent signals at the cost of including background, whereas narrower rings can miss some of the F-actin fluorescent surrounding the pillar. Sensitivity analysis with various ring widths (∼5.55µm 2, ∼8.33µm 2, ∼9.86µm 2, ∼10.41µm 2) ensured that the pillar-to-pillar correlations were not sensitive to the ring’s widths (Fig. S5). In addition we defined two negative control rings (of size ∼0.69µm 2 and ∼1.25µm 2) that were entirely placed within pillar voids that showed much weaker correlations (Fig. S5).

#### Topological distance

For each pillar, we identify the eight closest pillars as its immediate neighbors, forming the first level of topological distance (adjacent neighbors). The second level is defined by pillars that can be reached by two hops (two pillars away) from the central pillar. Subsequent levels are determined similarly, increasing the number of hops required to reach the pillars. Pillars classified under the second level and beyond are considered non-adjacent to the original pillar. See Fig. 1E.

### Cross-correlation analysis

For each cell at every topological distance, we computed the cross-correlation with a maximum lag of 3 frames (time lags from -3 to +3 frames, ± 90 seconds) for each pillar pair. To determine the correlation gain, we subtracted the maximum correlation within the lag range (-3 to +3) from the correlation at lag 0 (no time shift) (Fig. 1G). The correlation gain for a cell at a given topological distance is the average correlation gain across all pillars associated with that cell within the distance.

### Statistical analysis

Pearson correlation was used to measure the correlation between pillar pairs’ time series’.

Wilcoxon signed-rank statistical test was used to test whether the median of the correlation gain differences is significantly different from zero in Fig. 1H. The non-parametric Wilcoxon signed-rank test was chosen due to the small sample size and due to the unknown underlying distribution of our data. T-test was used to (1) compare adjacent pairs correlation to non adjacent pairs in both pillars height 5.3μm and 13.2μm in Fig. 2A, (2) the difference between the matched correlations of adjacent and nonadjacent pillar pairs distribution of 5.3μm versus 13.2μm height in Fig. 2A’s inset, (3) cell’s Δcorrelations of adjacent versus non-adjacent pillar pairs before and after the blebbistatin in both heights 5.3μm and 13.2μm in Fig. 2B, and (4) the difference between 5.3μm distribution and 13.2μm distribution of matched Δcorrelations before and after adding blebbistatin in Fig. 2B inset. T-test was chosen due to its effectiveness in comparing group means under the assumptions of normal distribution and equal variances. All significance tests were carried out with an α-value of 0.05, considering *P < 0.05, **P < 0.01, ***P < 0.001, ****P < 0.0001.

### Code and data availability

We are currently organizing our source code and processed data. We will make both publically available as soon as possible (before journal publications).

## Supplemental video legend

**Video S1**. 2D time-lapse confocal microscopy imaging of mouse embryonic fibroblasts (MEFs) expressing tdTomato-tractin positioned on top of an array of micropillars of 5.3-μm high and 2-μm wide, with a 4-μm distance between any two adjacent pillars. Timestamp is measured in minutes.

## Supplemental table legends

**Table S1**. Data description. Each row refers to a bulk of datasets including pillars height, cell type, treatment, temporal resolution, number of frames, number of cells, number of movies extracted from the cells and resolution.

## Funding and Acknowledgments

This research was supported by the Israeli Council for Higher Education (CHE) via the Data Science Research Center, Ben-Gurion University of the Negev, Israel (to AZ), by the Israel Science Foundation (ISF) grant (2516/21), and by the Wellcome Leap Delta Tissue program (to AZ). We thank Esraa Nsasra for critically reading the manuscript. HW is an incumbent of the David and Inez Myers Career Advancement Chair in Life Sciences. This research was supported by an Israel Science Foundation grant (1738/17) (HW) and the Rappaport Family Foundation (HW).

## Author Contribution

AZ and HW conceived the study. YG designed the experimental assay and performed all experiments. SH developed analytic tools, analyzed, and interpreted the data. AZ and HW mentored SH and YG. All authors drafted and edited the manuscript and approved its content.

## Supplementary information

**Figure S1.**
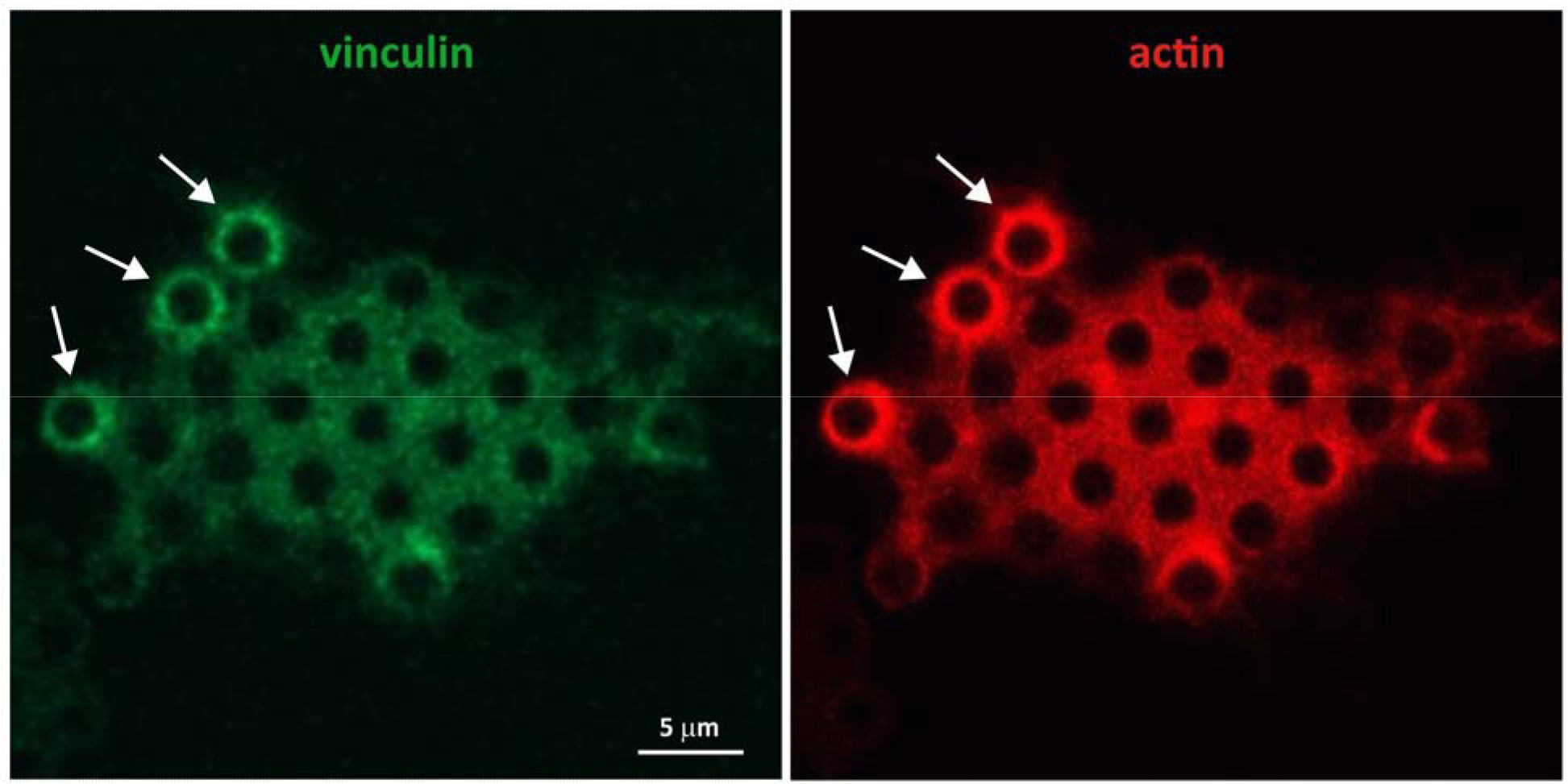
Adhesions form around the pillars. Example of a cell plated on 2 μm diameter pillars stained for vinculin and F-actin. Vinculin adhesions form around the pillars, evident by the ring pattern, which correspond to the polymerization of actin around the pillars (arrows).

**Figure S2.**
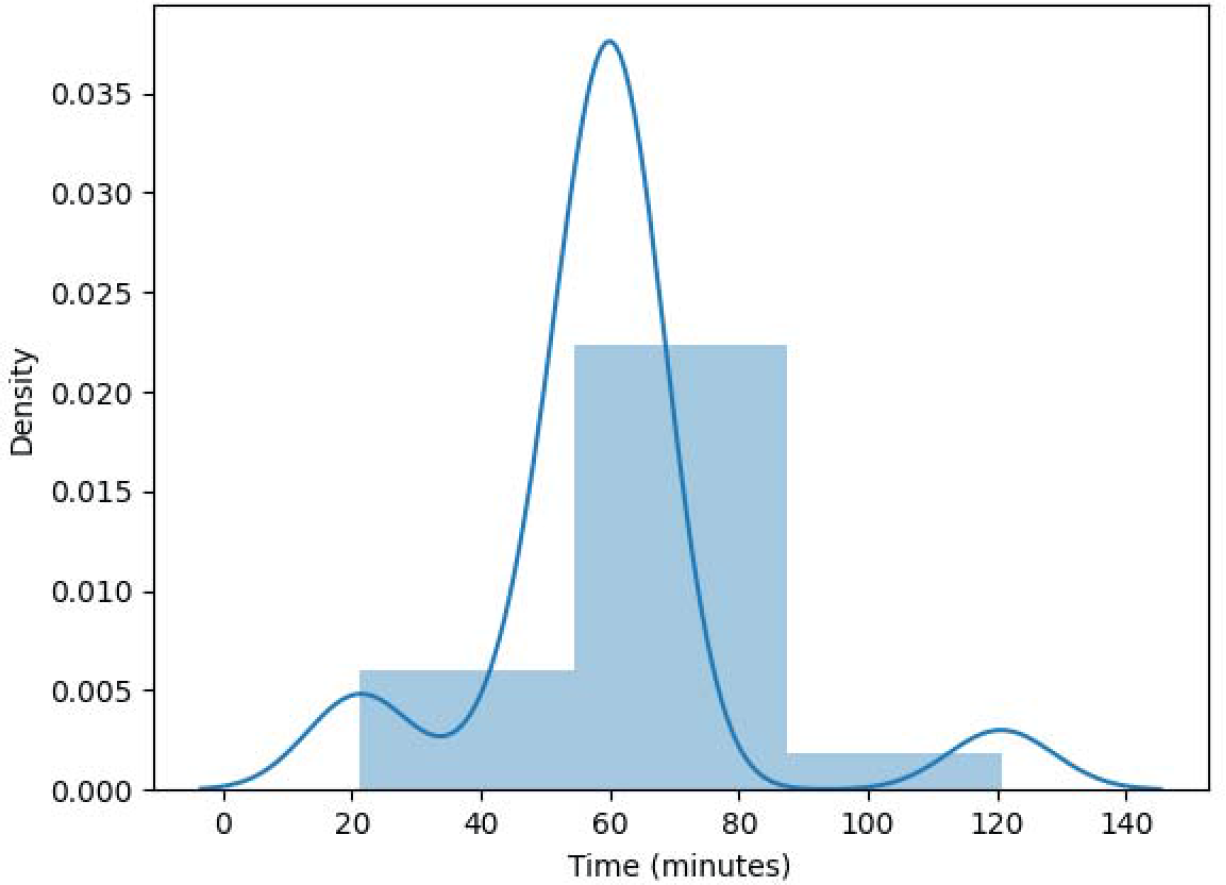
Distribution of the imaging duration for cells positioned on micropillars of height 5.3μm.

**Figure S3.**
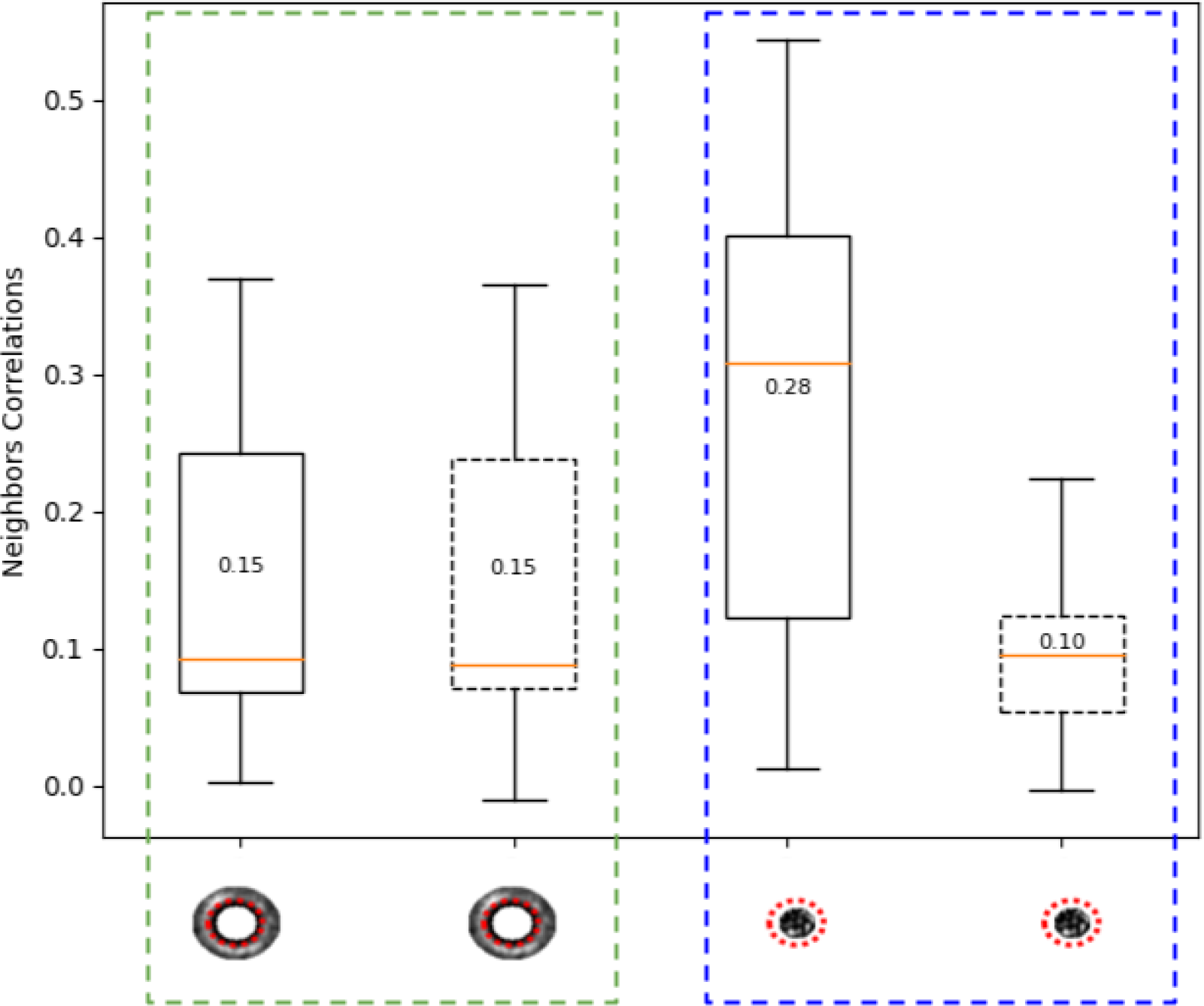
Control analysis: correcting for high background correlations through background normalization. Correlation in the control ring (highlighted in blue), which includes pillar voids, significantly decreases from an average of 0.28 to 0.1 after background normalization. In contrast, the original quantification ring (highlighted in green) maintains a consistent correlation value (average of 0.15) before and after background normalization.

**Figure S4.**
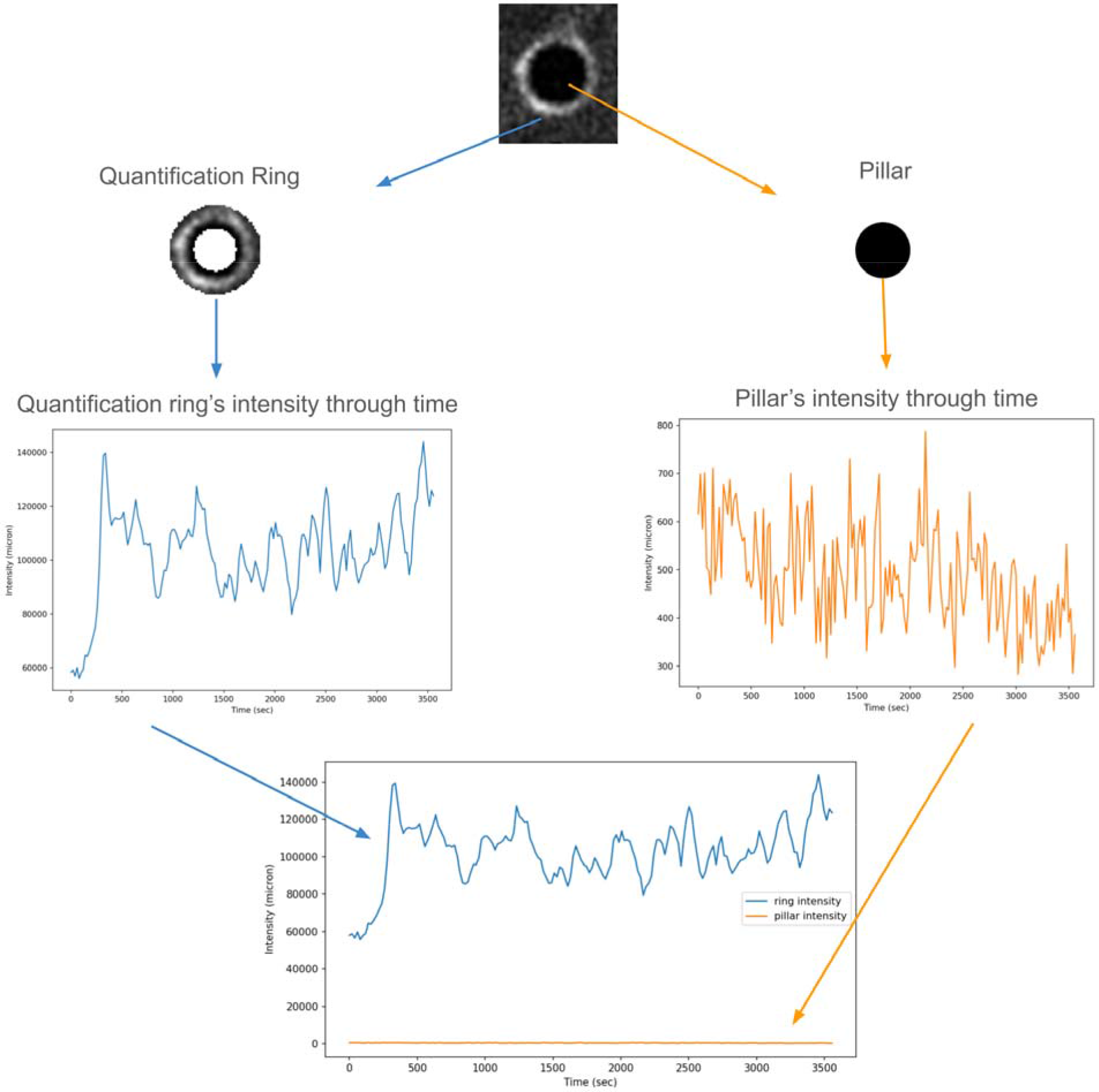
Quantification ring normalization. The pillar’s background was determined according to the intensity of the fluorescent signal within the pillar (orange) and was used to normalize the intensity of the fluorescent signal in the quantification ring (blue). The time series of each quantification ring was normalized by subtracting the mean intensity of all cells’ pillar voids at the corresponding time.

**Figure S5.**
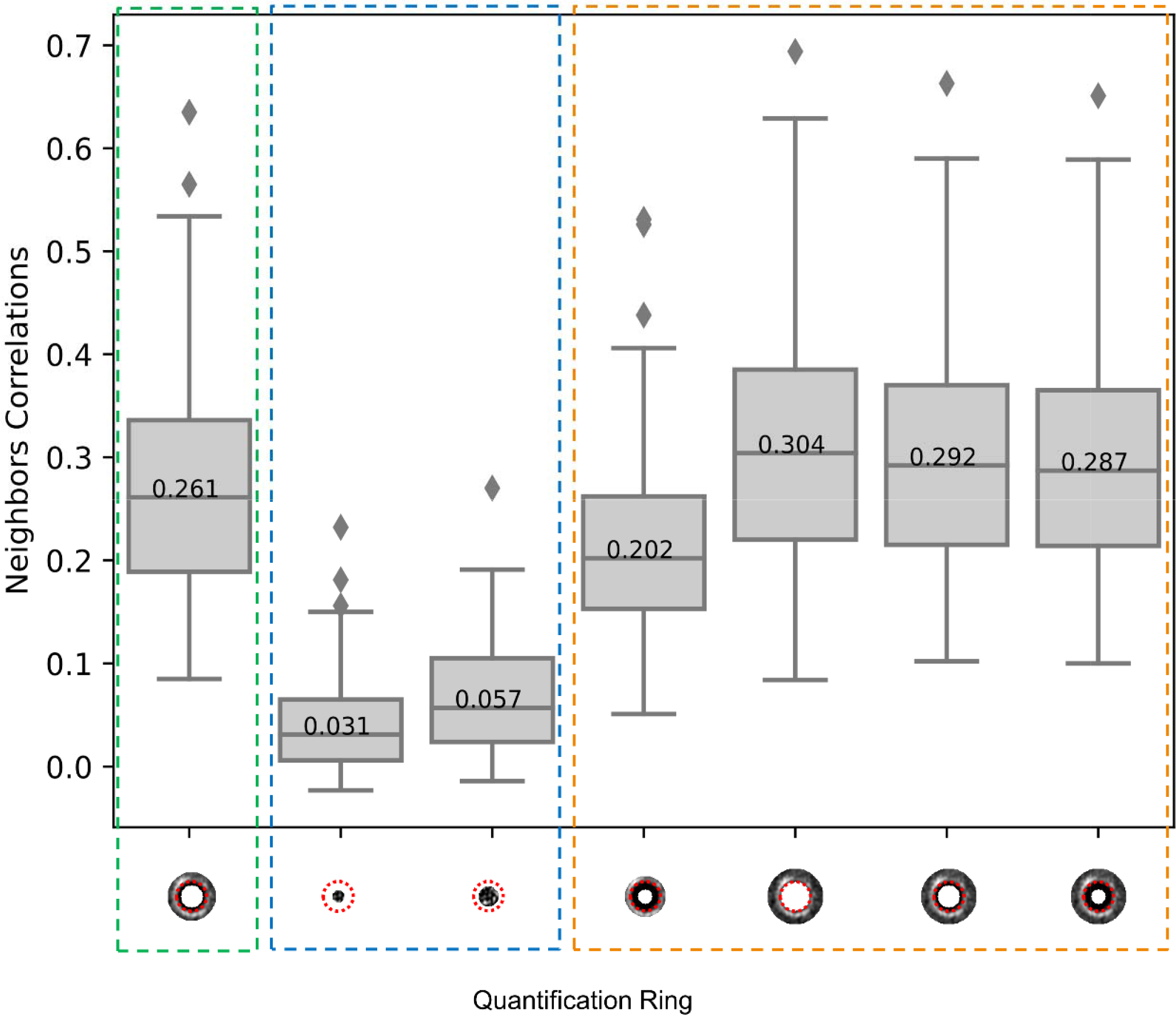
Sensitivity to quantification ring location and radius. Distribution of the correlation between adjacent pairs of pillars’ time-series for different quantification rings location and radii. X-axis: The pillar is marked with a red dashed line, and the quantification ring (gray-scale signal) is shown in respect to the pillar (left-to-right): ∼8.13, (quantification ring width). Dashed green rectangle indicates the quantification ring used in this study. Dashed blue rectangle indicates quantification rings within the pillar (negative control). Dashed orange rectangle indicates quantification rings with varying sizes (sensitivity analysis). This analysis indicates that pillar-to-pillar synchronization must be measured externally to the pillar and is not very sensitive to the radius of the quantification ring.

**Figure S6.**
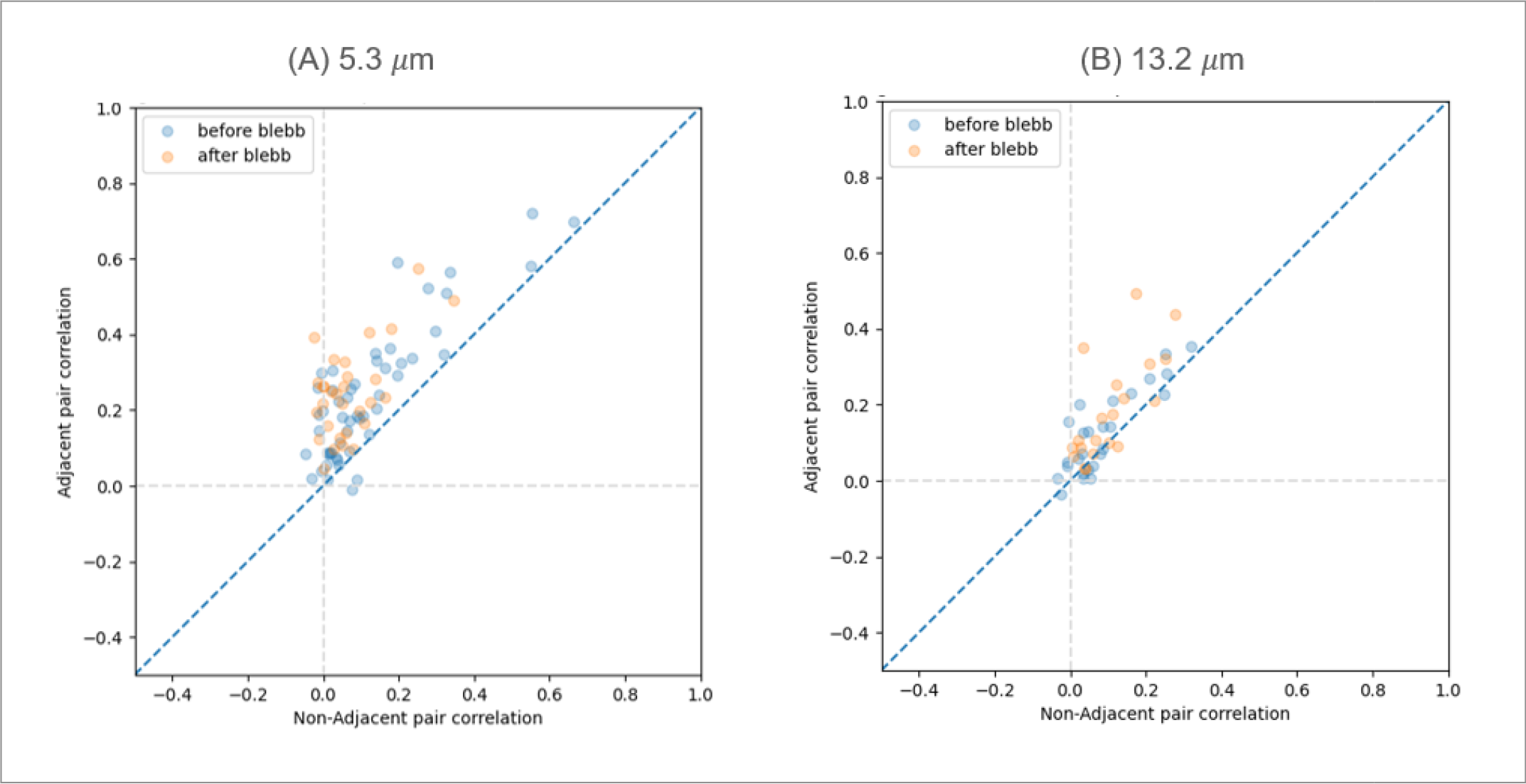
Adjacent-versus-non-adjacents pair analysis before and after adding blebbistatin. Each data point records the mean correlation of a cell’s adjacent and non-adjacent pillar pairs of an experiment. (1) Cells on top of pillars of height 5.3 m. Before adding blebbistatin (N = 52, blue), adjacent pairs had higher correlations than non-adjacent pairs in 96.15% (50/52) of cells. After adding blebbistatin (N = 30, orange), adjacent pairs had higher correlations than non-adjacent pairs in all 30 cells. (2) Cells on top of pillars of height 13.2 m. Before adding blebbistatin (N = 26, blue), adjacent pairs had higher correlations than non-adjacent pairs in 65.38% (17/26) of cells. After adding blebbistatin (N = 20, orange), adjacent pairs had higher correlations than non-adjacent pairs 75% (15/20) of cells.

